# Binary-SPA: A Reference-Free Method for Cell Annotation in High-Resolution Spatial Transcriptomics

**DOI:** 10.64898/2026.03.17.712369

**Authors:** Honghao Bi, Wenjie Cai, Pan Wang, Kehan Ren, Inci Aydemir, Ermin Li, Johanna Melo-Cardenas, Matthew J. Schipma, Ching Man Wai, Peng Ji

## Abstract

Accurate cell type annotation remains a major challenge in high-resolution spatial transcriptomics analysis. Current approaches primarily rely on label transfer from single-cell RNA sequencing (scRNA-seq) reference datasets or marker-based annotation of transcriptionally defined clusters. While these methods are widely used, they have critical limitations. Label transfer approaches depend on the availability of a well-matched scRNA-seq reference. Marker-based annotation methods often suffer from accuracy and limited coverage. To address these challenges, we developed Binary-SPA, a computational framework for cell-type annotation of high-resolution spatial transcriptomics data. Binary-SPA performs annotation in two stages. First, a binary classification step identifies high-confidence cells using predefined marker sets. These confidently annotated cells are then used as an internal reference for anchor-based label transfer in the second stage. Binary-SPA outperforms conventional marker-based annotation approaches and existing label transfer methods across multiple high-resolution spatial transcriptomics platforms, preservation methods, and tissue types. While label transfer methods achieve high accuracy only when same-tissue scRNA-seq references are available, and decline substantially when relying on independent datasets, Binary-SPA matches this performance with 100% annotation coverage while eliminating the need for external reference data entirely. Binary-SPA demonstrates robust performance even in challenging specimens such as bone marrow biopsies, and validation against matched COMET protein expression data confirmed strong concordance between transcriptomic- and protein-based cell identities. Binary-SPA thus provides a robust, reference-free solution for spatial transcriptomics annotation with broad applicability to research and clinical specimens.

## Introduction

Spatial transcriptomics has emerged as a powerful technology for elucidating cell-cell interactions and functional niches within a spatial context^1–4^. To fully leverage these insights, accurate and robust cell annotation is essential for downstream analyses. However, existing annotation methods often struggle to accurately identify all cell types. A common workflow is to cluster cells and then examine the expression of known marker genes across clusters^5,6^. Clusters are then annotated based on the highly expressed markers. In practice, however, this approach often yields conflicting marker profiles: a single cluster may express markers associated with multiple cell types. These inconsistencies arise because unsupervised clustering and traditional cell typing rely on fundamentally different principles: while the traditional method uses a small set of marker proteins to distinguish cell populations, clustering algorithms classify cells based on global RNA expression patterns. To address this mismatch, many studies increase clustering resolution to generate more clusters. As a result, the clustering-based analyses often yield overly detailed annotations and/or improvised cell types.

Label transfer annotation methods provide powerful tools for cell type identification^7,8^. However, their performance depends heavily on the reference dataset, which is typically a scRNA-seq reference annotated using the clustering-based methods described above. Consequently, any inaccuracies in the original reference can propagate into downstream analyses. Moreover, inter-individual variation poses a significant challenge for label transfer. For example, cells from diseased tissues often undergo global transcriptomic changes, which can shift their expression profiles away from those observed in healthy reference datasets and lead to misannotation^9,10^. Although some studies use matched scRNA-seq data from the same tissue as a reference, this approach is often not feasible, particularly for archived clinical samples. As a result, many studies rely on “similar” samples as a reference, which can compromise annotation accuracy.

To improve the annotation accuracy in high-resolution spatial transcriptomics, we developed Binary-SPA (Binary Self-referenced Projection Annotation). Benchmarking results show that Binary-SPA achieves performance comparable to that of label transfer methods when using the same tissue as the reference data. Importantly, when the reference dataset is derived from independent studies, Binary-SPA outperforms all tested methods. Binary-SPA performs cell-by-cell annotation based on marker genes and achieves a 100% annotation rate with high accuracy. It also allows annotation with a user-defined set of cell types, ensuring results are better aligned with traditional cell type frameworks and easier to interpret biologically.

## Results

### Conceptualization of Binary-SPA for high-resolution spatial transcriptomics analysis

Binary-SPA consists of two steps: a marker-based binary annotation step (Binary) and a self-referenced projection annotation step (SPA) (**Fig. 1**). In the Binary step, cells expressing predefined marker genes are identified and annotated with high confidence. These confidently annotated cells are then used as a reference dataset for anchor-based annotation of the remaining cells in the SPA step. We reasoned that although some cells may lose detectable marker expression at the RNA level, they should still retain transcriptomic profiles characteristic of their true cell type.

**Fig. 1.**
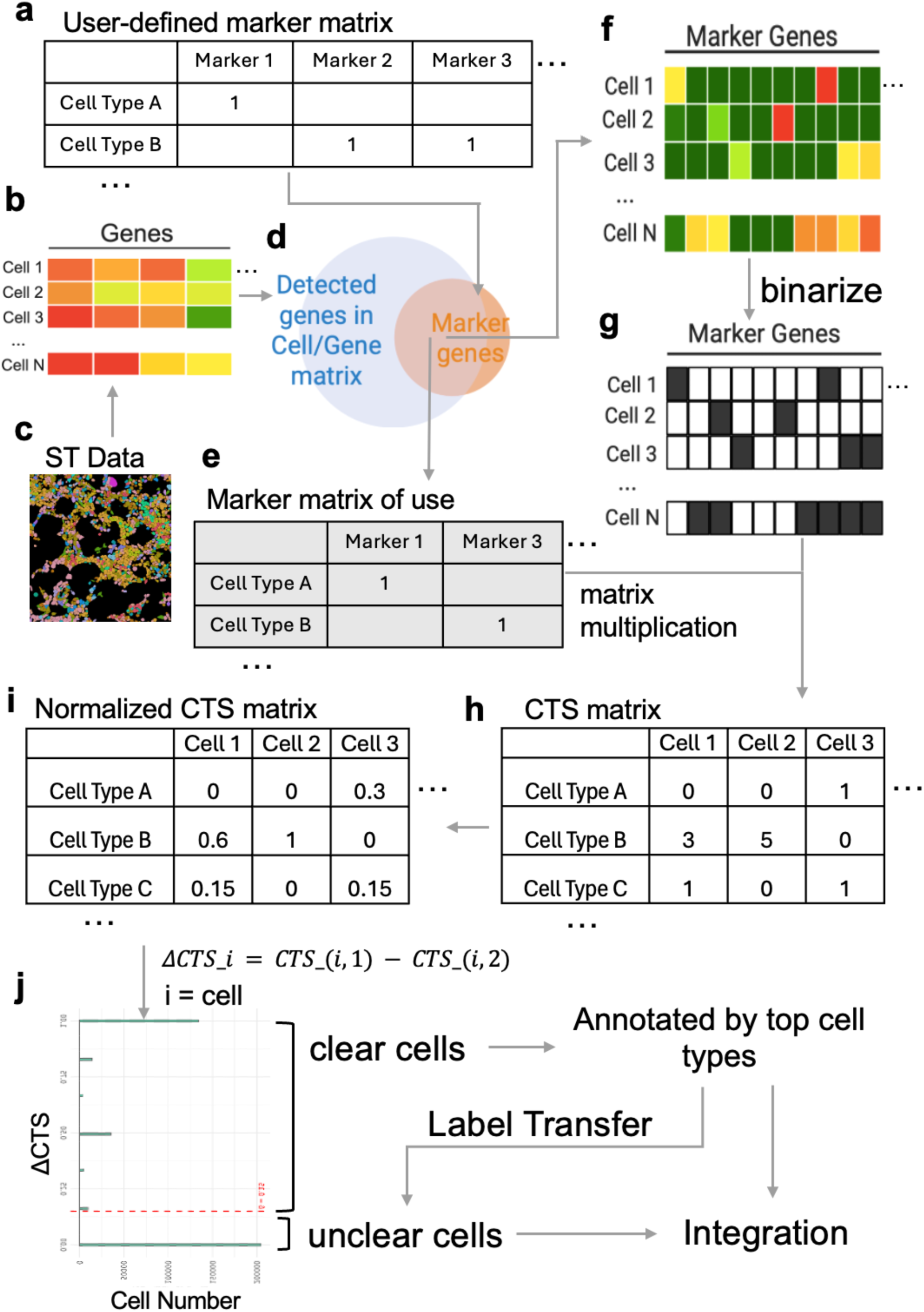
Overview of the Binary-SPA workflow. **a**, User-defined marker matrix with cell types as rows and marker genes as columns. Markers expected to be positively expressed in each cell type are assigned a value of 1. All other entries are set to 0. **b-c**, Cell-by-gene expression matrix (b) derived from spatial transcriptomics (ST) data (c), where rows represent cells, columns represent genes, and color intensity indicates expression level. **d-e**, Genes shared between the expression matrix and the user-defined marker matrix were identified (d) and used to construct a platform-specific marker matrix of use, retaining only marker genes present in the spatial dataset (e). **f**, Cell-by-marker gene expression matrix, including only overlapping marker genes. **g**, Binarized marker expression matrix, where detectable expression is encoded as 1 (black) and non-detectable expression as 0 (white). **h**, Cell type score (CTS) matrix generated by matrix multiplication of the marker matrix of use and the transposed binarized expression matrix. Each entry represents the number of detected markers for a given cell type in a given cell. **i**, Normalized CTS matrix following min-max scaling by cell type for cross-cell-type comparisons. **j**, ΔCTS calculation and cell classification. For each cell, ΔCTS is computed as the difference between the highest and second-highest normalized CTS values. Cells with ΔCTS exceeding a user-defined threshold (dashed red line) are classified as clear cells and annotated by their top-scoring cell type. Unclear cells receive annotations via anchor-based label transfer, using clear cells as the internal reference, resulting in complete annotations for all cells.

The Binary step begins with quality control, in which unsupervised clustering is performed to verify that no unexpected cell populations are present in the testing sample. Differentially expressed genes are examined across clusters to identify previously unannotated cell types resulting from disease or other biological processes, or cells originating from other tissues (e.g., metastatic populations). After quality control, a user-defined marker matrix is generated with cell types as rows and marker genes as columns. Marker genes expected to be positively expressed in each cell type, based on prior knowledge, are assigned a value of 1, while all other entries are set to 0 (**Fig. 1a**; exemplified in **Extended Data Tables 1, 2**). If unexpected populations were identified during quality control, corresponding markers are added to this matrix.

Next, a cell-by-gene expression matrix is generated from the spatial transcriptomics data, where each row represents a cell, each column represents a gene, and each entry contains the expression level of that gene in that cell (**Fig. 1b, c**). Because high-resolution spatial transcriptomics platforms vary in their gene coverage, for example, Xenium 5k detects approximately 5,000 genes while other platforms may detect more or fewer, some marker genes in the user-defined matrix may not be present in the dataset. To ensure compatibility, a platform-specific marker matrix of use is generated: marker genes absent from the spatial dataset are removed from the user-defined matrix, retaining only those that can be evaluated (**Fig. 1d, e**). The cell-by-gene expression matrix is subsequently subset to include only these overlapping marker genes (**Fig. 1f**).

This subset matrix is then binarized: detectable expression is assigned a value of 1 (black), and non-detectable expression is assigned a value of 0 (white) (**Fig. 1g**). This binarization converts marker gene expression into a simple on/off signal, prioritizing the presence of multiple markers over the magnitude of expression for any single marker, an approach that more closely mirrors classical immunophenotyping logic.

The cell type score (CTS) is then calculated for each cell by counting the number of detected markers corresponding to each cell type. This is achieved through matrix multiplication of the marker matrix of use and the transposed binarized expression matrix, generating a CTS matrix in which each cell receives a score for each predefined cell type (**Fig. 1h**). The CTS matrix is then normalized by cell type using min-max scaling, such that CTS values range from 0 to 1 for each cell type, enabling cross-cell-type comparisons (**Fig. 1i**).

For each cell, ΔCTS is calculated as the difference between the highest and second-highest normalized CTS values (**Fig. 1j**). Cells with ΔCTS exceeding a user-defined threshold are designated “clear cells” and annotated according to their top-scoring cell type. The remaining “unclear cells” are initially labeled as NA.

In the SPA step, clear cells serve as the reference dataset for anchor-based label transfer onto unclear cells using the MapQuery function in Seurat. Because all cells originate from the same sample and are subject to identical experimental and biological conditions, batch effects between clear and unclear cells are minimized, and unclear cells are expected to share transcriptomic signatures with clear cells of the same identity, which enables accurate label transfer. The predicted annotations are merged back into the original dataset, yielding a complete annotation of all cells. We term this self-referencing strategy Self-Projection Annotation (SPA) (**Fig. 1j**).

### Validation of Binary-SPA in spatial transcriptomics data

To evaluate the performance of Binary-SPA, we applied the method to Xenium 5k datasets from a published benchmarking study for high-resolution subcellular spatial transcriptomics^11^, in which cell type annotations were independently established using both transcriptomic and proteomic references. In this study, scRNA-seq reference data were generated from the same fresh-frozen tissue used for the spatial profiling, and cell types were assigned to Xenium data using a voting-based strategy that integrated five independent annotation methods. These annotations were further validated using CODEX protein imaging data generated from adjacent tissue sections, providing an orthogonal benchmark for cell type identification. Three tumor types were analyzed: colon adenocarcinoma (COAD), hepatocellular carcinoma (HCC), and ovarian cancer (OV). For Binary-SPA analysis, we constructed two user-defined marker matrices: one shared between the COAD and OV datasets and a second HCC-specific set that included hepatocyte markers (**Extended Data Tables 1, 2**). This experimental design allowed us to directly compare Binary-SPA’s self-referencing approach against methods that had access to matched scRNA-seq data from the same tissue.

Binary-SPA successfully annotated all cells across the three datasets. Spatial cell type distributions recapitulated tissue architecture visible in corresponding H&E-stained sections, with epithelial tumor regions, stromal compartments, and immune infiltrates appropriately localized in the Binary-SPA annotation maps (**Extended Data Fig. 1a-f**). Notably, Binary-SPA showed a higher annotation rate than the reported voting-based method, which was designed to maximize annotation accuracy through multi-method consensus (**Fig. 2a, b**). Quantitative comparison of annotation coverage revealed that Binary-SPA achieved 100% annotation rate across all samples, compared to approximately 90% for the original voting-based approach that relied on matched scRNA-seq references (**Fig. 2a**). The unannotated cells in the original study likely represent cells with ambiguous transcriptomic profiles that failed to reach consensus across the five reference-dependent methods. Binary-SPA’s self-referencing label transfer strategy successfully assigned identities to these challenging cells (**Extended Data Fig. 2**).

**Fig. 2.**
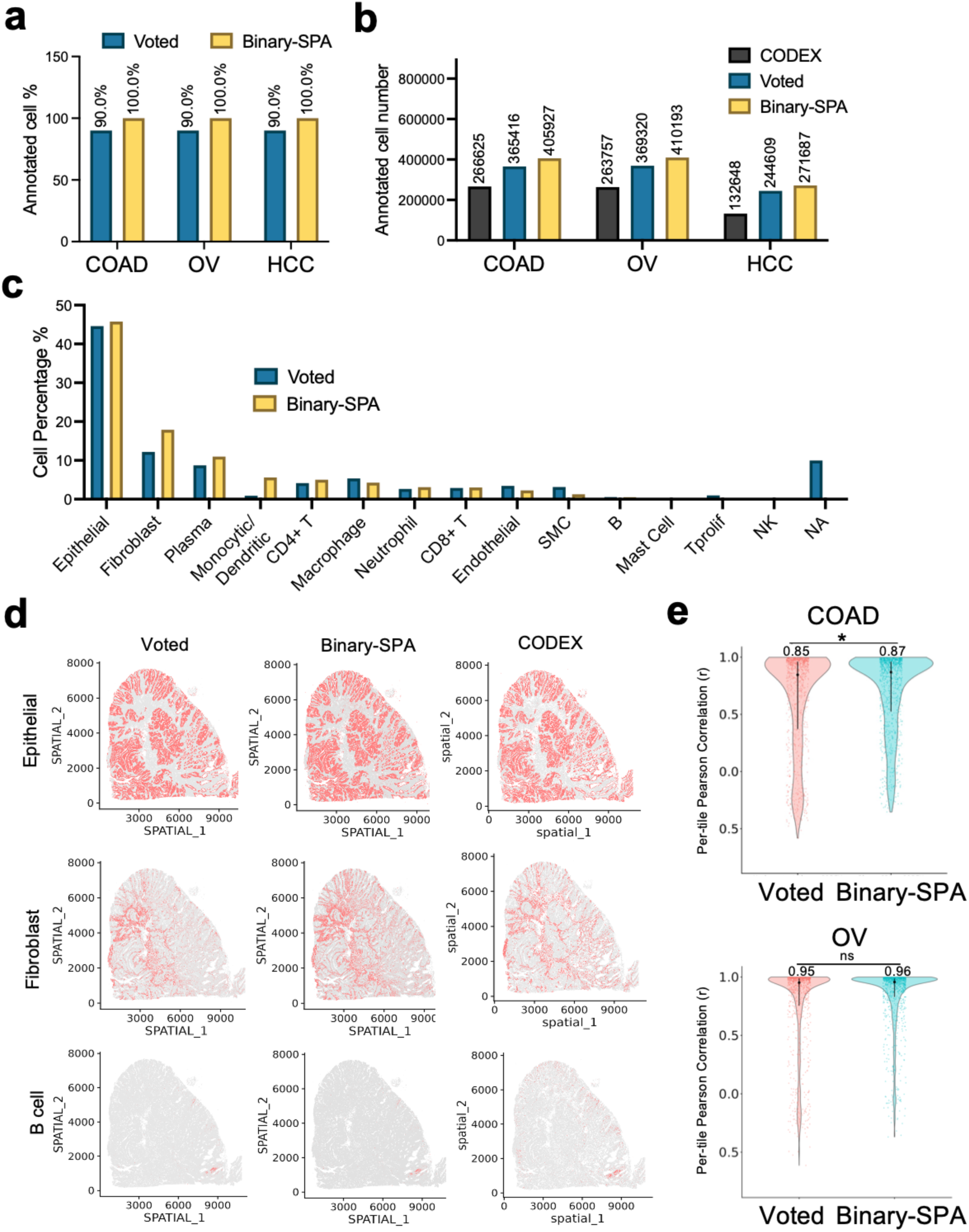
Annotation of Xenium data using Binary-SPA. **a**, Comparison of annotation coverage between the voting-based method (Voted) and Binary-SPA across colon adenocarcinoma (COAD), ovarian cancer (OV), and hepatocellular carcinoma (HCC) Xenium datasets. Percentages indicate the proportion of total cells successfully annotated by each method. **b**, Absolute number of annotated cells for each method. CODEX protein-based annotations serve as the reference standard. Voted and Binary-SPA annotations are shown for comparison. **c**, Cell type composition comparison between Voted and Binary-SPA methods in the COAD dataset. Bar heights represent the percentage of each cell type among all annotated cells. **d**, Spatial dimplots illustrating representative cell types (epithelial cells, fibroblasts, and B cells) across Voted, Binary-SPA, and CODEX annotations in the COAD dataset. **e**, Per-tile Pearson correlation analysis comparing Voted and Binary-SPA annotations against CODEX ground truth in COAD (top) and OV (bottom) datasets. Spatial transcriptomics data were divided into 1,000 tiles, and cell type proportions within each tile were correlated with matched CODEX measurements. Violin plots show the distribution of correlation coefficients across all tiles; median r values are indicated above each violin. Center dots indicate medians; vertical lines represent 95% confidence intervals. Statistical significance assessed by the Wilcoxon test; * p < 0.05, ns = not significant.

Cell type proportion analysis demonstrated strong concordance between Binary-SPA and the original voting-based annotations (**Fig. 2c**). Across all three datasets, the relative abundances of major cell populations, including epithelial cells, fibroblasts, plasma cells, and immune subsets, were highly correlated between methods. To further validate annotation accuracy, we compared spatial distribution patterns of individual cell types across Binary-SPA, the original voting strategy, and CODEX protein-based annotations (**Fig. 2d**). Representative cell types, such as epithelial cells, fibroblasts, and B cells, showed comparable spatial localization across all three methods. Notably, Binary-SPA fibroblast annotations more closely matched CODEX protein-based detection than the voting-based approach, which appeared to underestimate fibroblast density in certain regions.

To further validate Binary-SPA and directly benchmark its performance quantitatively, we performed a tiled Pearson correlation analysis. Each spatial map was partitioned into 1,000 tiles, and within each tile, cell-type proportions derived from the reported voted annotations and from Binary-SPA were independently compared with CODEX annotations. Per-tile Pearson correlation coefficients (r) were then aggregated for downstream statistical analysis. We found that both annotation strategies exhibited strong concordance with CODEX, with the majority of tiles showing correlation values approaching 1. In the COAD dataset, Binary-SPA achieved a significantly higher median correlation than the voted annotations (r = 0.87 vs. 0.85; Wilcoxon test, p = 0.042) (**Fig. 2e**). In the OV dataset, Binary-SPA showed a comparable median correlation (r = 0.96 vs. 0.95; p = 0.175). Tile-based analysis was not performed for HCC due to insufficient CODEX data quality. Collectively, these data show that Binary-SPA outperforms a consensus voting strategy that integrates five annotation methods with matched scRNA-seq references, while requiring no external reference data.

### Benchmarking Binary-SPA against reference-dependent and marker-based annotation methods

Because most studies rely on a single annotation method rather than consensus voting, benchmarking against individual methods provides a more realistic assessment of practical performance. We therefore benchmarked Binary-SPA against seven individual methods used for high-resolution spatial transcriptomics in the original study: five reference-dependent label-transfer methods (SELINA^12^, SPOINT^13^, TACCO^14^, CellTypist^15^, and Tangram^16^) and two marker-based methods (TACIT^17^ and ScType^18^). Among reference-dependent methods, substantial performance variability was observed (**Fig. 3a**, **Extended Data Fig. 3**). Tangram showed marked deviations from the consensus, with pronounced overestimation of certain populations. TACCO underestimated epithelial cells in COAD and hepatocytes in HCC while overestimating macrophages. SPOINT achieved only ∼90% annotation coverage. SELINA, despite fine-tuning with matched reference data, exhibited significantly lower annotation rates. Only CellTypist achieved performance comparable to Binary-SPA, but again, required matched scRNA-seq reference data that is often unavailable in practice. In contrast, Binary-SPA achieved cell type proportions highly concordant with the consensus benchmark across all three tumor types without requiring any external reference.

**Fig. 3.**
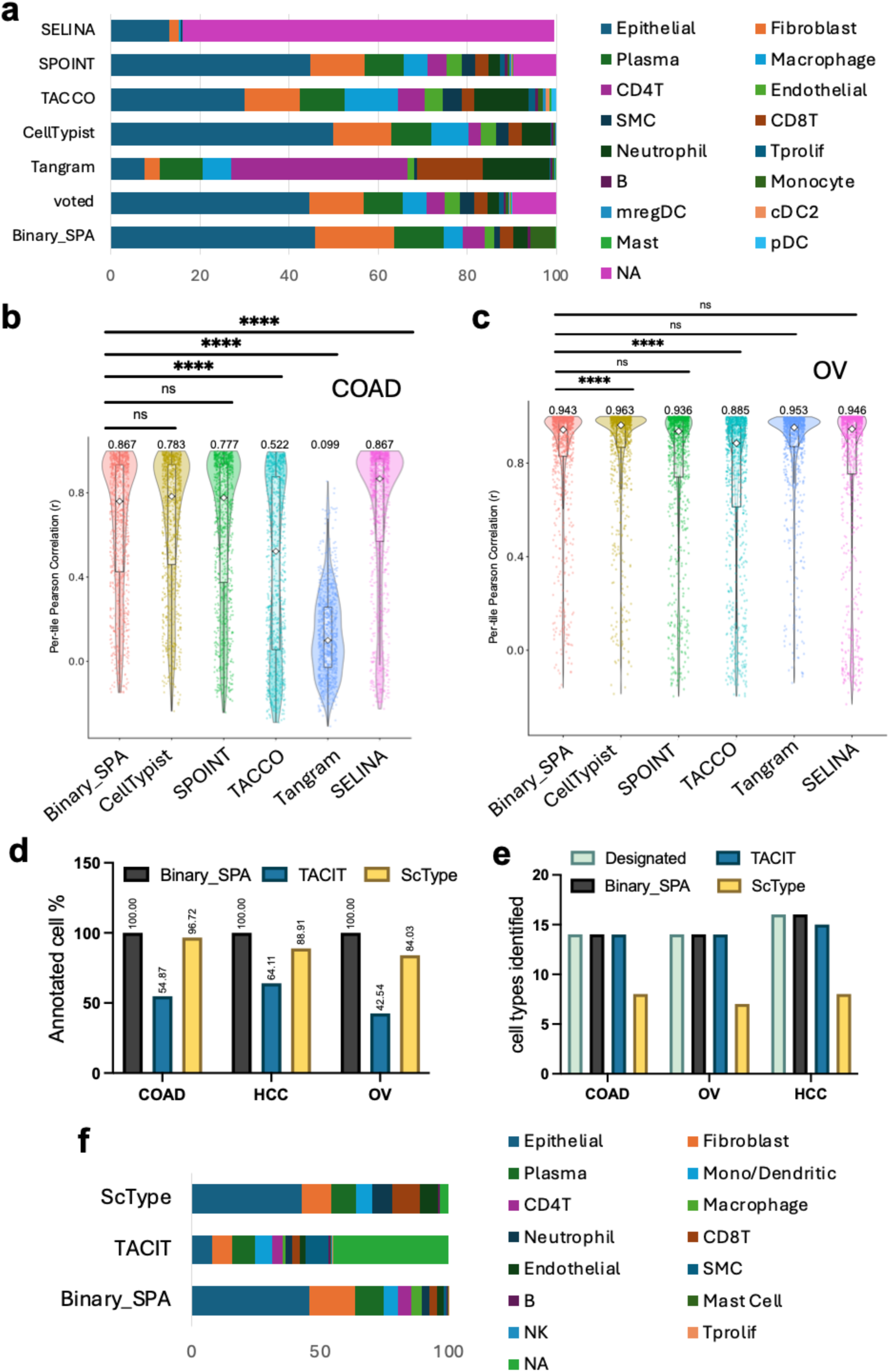
Benchmarking Binary-SPA against other annotation methods. **a**, Cell type composition across seven annotation methods (SELINA, SPOINT, TACCO, CellTypist, Tangram, Voted, and Binary-SPA) in the COAD dataset. Stacked bars represent the proportion of each cell type among all annotated cells. **b**, Per-tile Pearson correlation analysis comparing each annotation method against CODEX ground truth in the COAD dataset. Spatial transcriptomics data were divided into 1,000 tiles, and cell type proportions within each tile were correlated with matched CODEX measurements. Violin plots show the distribution of correlation coefficients across tiles; median r values are indicated above each violin. Statistical significance compared to Binary-SPA was assessed by the Wilcoxon test; **** p < 0.0001, ns = not significant. **c**, Per-tile Pearson correlation analysis in the OV dataset, presented as in (b). **d**, Annotation coverage comparison among reference-free methods (Binary-SPA, TACIT, and ScType) across COAD, HCC, and OV datasets. Percentages indicate the proportion of total cells successfully annotated. **e**, Number of cell types identified by each indicated method compared to the predefined cell type panel. Gray bars indicate predefined cell types; colored bars indicate detected cell types. **f**, Cell type composition comparison among Binary-SPA, TACIT, and ScType in the COAD dataset.

To quantitatively compare spatial accuracy, we used CODEX protein data as ground truth and the tiled Pearson correlation analysis described above. Binary-SPA achieved consistently high concordance with CODEX across datasets, whereas Tangram and TACCO showed substantially lower correlations in COAD, and TACCO underperformed in OV (**Fig. 3b, c**). While SELINA exhibited high accuracy, its low annotation coverage limits practical utility (**Extended Data Fig. 3a-c**).

Binary-SPA also substantially outperformed marker-based methods. Using identical marker gene sets (**Extended Data Table 3**), TACIT achieved only 42-65% annotation coverage across datasets (**Fig. 3d**). ScType achieved higher coverage (84-100%) but failed to recover multiple cell types, identifying only 7-8 of 14 designated populations in COAD and OV (**Fig. 3e**). Binary-SPA achieved 100% coverage while recovering all designated cell types, with proportions closely matching the consensus benchmark (**Fig. 3d-f**; **Extended Data Fig. 4a, b**). Together, these results indicate that Binary-SPA delivers reference-level accuracy while preserving the flexibility of reference-free marker-based annotation.

### Binary-SPA enables robust annotation across high-resolution spatial transcriptomics platforms and sample preservation methods

A critical requirement for any annotation method is generalizability across different technological platforms and sample types. Visium HD represents a distinct high-resolution spatial transcriptomics approach from Xenium, utilizing sequencing-based capture rather than imaging-based detection, with different gene panels, sensitivity profiles, and spatial resolution characteristics. Furthermore, clinical and research applications frequently require analysis of both fresh-frozen and formalin-fixed paraffin-embedded (FFPE) specimens, which exhibit markedly different RNA quality and expression profiles. We therefore evaluated whether Binary-SPA’s reference-free annotation framework could maintain performance across these technical and biological variables.

We applied Binary-SPA to COAD Visium HD datasets from the same benchmarking study, analyzing both FFPE tissue from sections adjacent to the Xenium analysis and fresh-frozen tissue from the same tumor specimen. Analysis was performed using 8 µm bins, which aggregate transcripts into spatial units approximating single-cell resolution, consistent with the original study. Using this bin size, Binary-SPA successfully reconstructed the spatial architecture of the tumor microenvironment in both sample types (**Fig. 4a-d, Extended Data Table 4**). Epithelial tumor nests, surrounding stromal fibroblasts, and infiltrating immune populations were appropriately localized and corresponded to histological features visible in matched H&E sections (**Fig. 4b, d**). Notably, during the quality control step, a population exhibiting features of both smooth muscle cells and fibroblasts was identified, likely representing myofibroblasts or reflecting mixed transcripts within 8 µm bins; myofibroblasts were therefore added to the marker set for subsequent analyses.

**Fig. 4.**
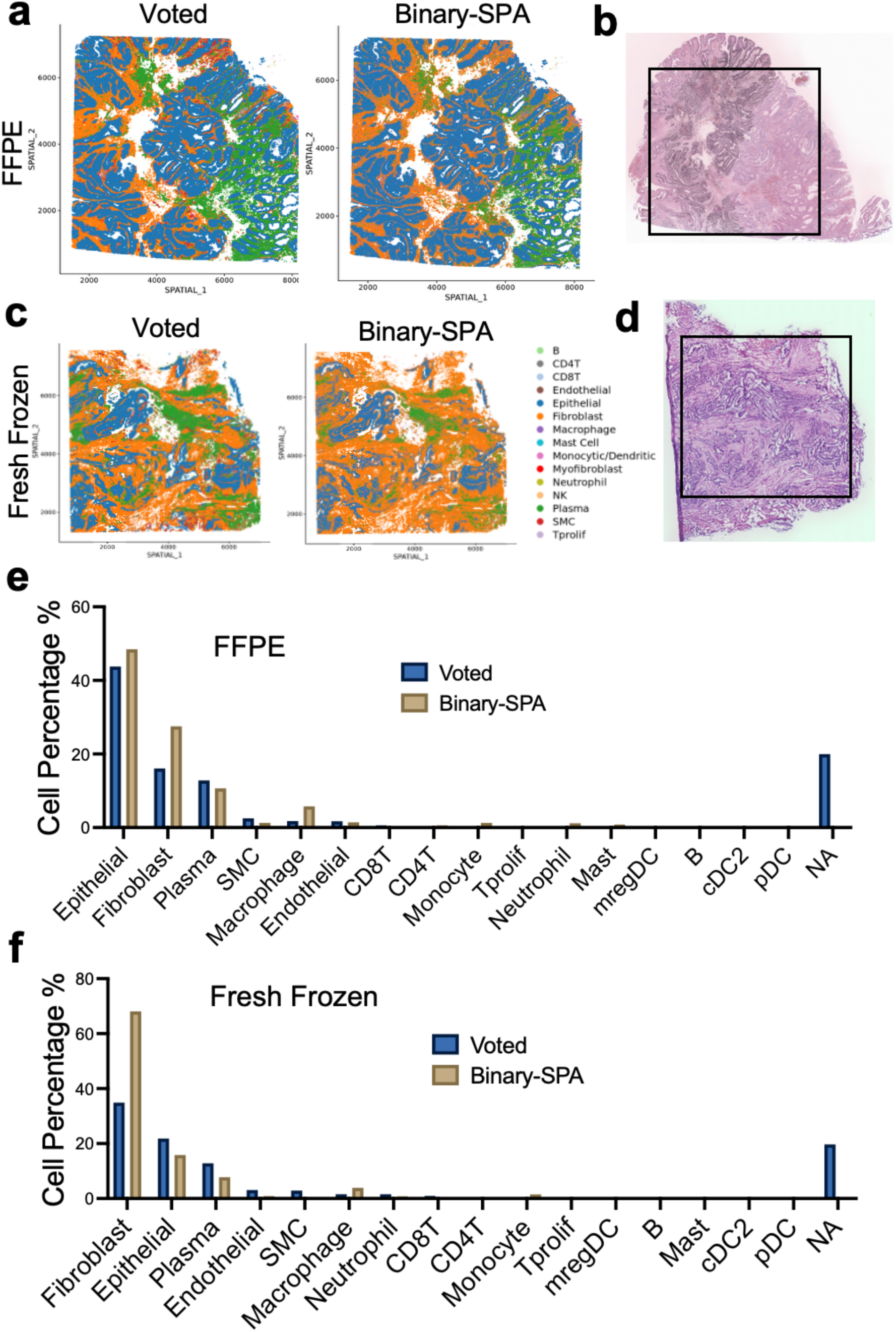
Binary-SPA performance across FFPE and fresh-frozen tissue preparations. **a**, Spatial distribution of cell type annotations by Voted and Binary-SPA methods in an FFPE COAD sample. Each dot represents a single cell colored by assigned cell type. Unannotated cells from the original study were filtered out to facilitate a direct comparison. **b**, Corresponding H&E-stained section of the FFPE sample shown in (a). The boxed region indicates the area of interest. **c**, Spatial distribution of cell type annotations by Voted and Binary-SPA methods in a fresh-frozen COAD sample. Unannotated cells from the original study were filtered out to facilitate a direct comparison. **d**, Corresponding H&E-stained section of the fresh-frozen sample shown in (c). The boxed region indicates the area of interest. **e-f**, Cell type composition comparison between Voted and Binary-SPA methods in the FFPE (e) and fresh frozen (f) COAD samples. Bar heights represent the percentage of each cell type among all annotated cells.

Binary-SPA required no modification when transitioning from Xenium to Visium HD. The same marker-based framework automatically adapts to different gene panels via an intersection step (**Fig. 1d**) that filters markers to platform-detected genes. Quantitative comparison revealed that Binary-SPA achieved 100% annotation coverage on both FFPE and fresh-frozen datasets, whereas the original voting-based approach left approximately 20% of spatial bins unannotated (**Fig. 4e, f**). Cell type proportions showed strong concordance between Binary-SPA and the original consensus annotations, with major populations including epithelial cells, fibroblasts, and plasma cells detected at comparable frequencies across both preservation conditions. These results demonstrate that Binary-SPA maintains accurate, complete annotation across platforms and sample types without requiring platform-specific optimization or external reference data.

### Binary-SPA captures clinically meaningful disease progression in bone marrow biopsies

Among tissues and biological systems, bone marrow presents particular challenges for spatial transcriptomics analysis. The hematopoietic system comprises large populations of closely related cell types that exhibit continuous developmental and functional states,^19–21^ resulting in transcriptomic continua rather than well-separated discrete clusters^22^. Additionally, bone marrow is a highly heterogeneous tissue composed primarily of nonadherent cells that are closely related yet functionally diverse. Furthermore, bone marrow sample processing typically requires chemical decalcification, which causes substantial RNA degradation and further complicates accurate spatial transcriptomic profiling.

To evaluate Binary-SPA in this challenging tissue context, we reanalyzed bone marrow Xenium datasets from a published study that employed an EDTA-based decalcification protocol to preserve RNA integrity^23^. The dataset comprises 21 Xenium 5k samples: 4 normal bone marrow controls, 2 from patients with monoclonal gammopathy of undetermined significance (MGUS), 5 from patients with smoldering multiple myeloma (SM), and 10 from patients with multiple myeloma (MM).

Binary-SPA successfully annotated cells and recapitulated spatial structures in these bone marrow core biopsy samples (**Fig. 5a, Extended Data Table 5**). Cell-type proportion analysis revealed a stepwise increase in plasma cell abundance from control to MGUS, SM, and MM samples, a progression consistent with the known disease biology of plasma cell neoplasms (**Fig. 5a**). To benchmark against reference-dependent approaches, we performed parallel annotation using SingleR^24^ with published bone marrow scRNA-seq reference data, following the original study design. SingleR failed to detect significant differences in plasma cell proportions between SM and control/MGUS groups, whereas Binary-SPA clearly identified this clinically meaningful progressive increase (**Fig. 5b, c**). These findings were further supported by plasma cell spatial distributions that showed concordance with CD138 immunohistochemistry in the original dataset (**Fig. 5d, e**).

**Fig. 5.**
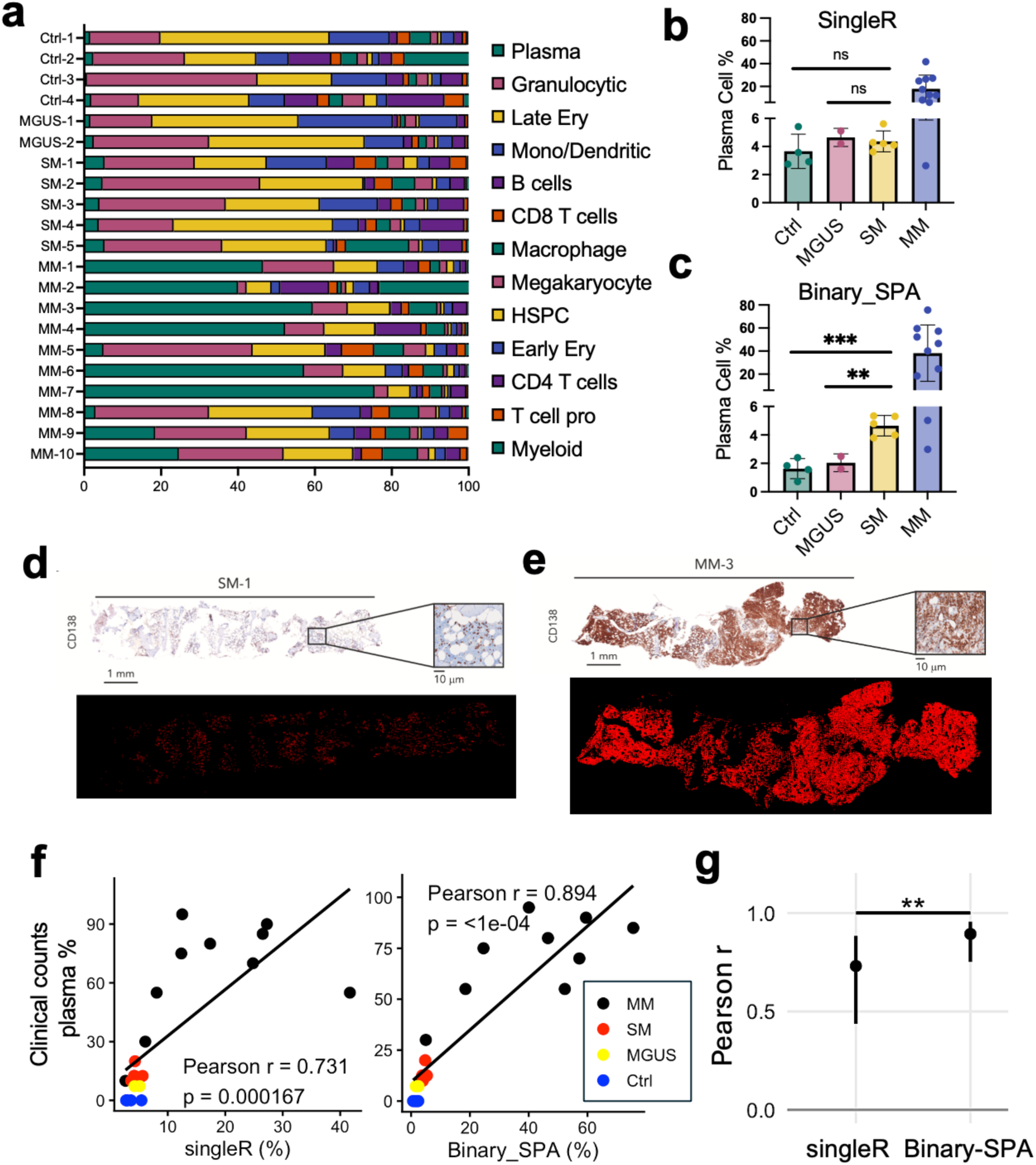
Binary-SPA validation in bone marrow spatial transcriptomics with clinical correlation. **a,** Cell type composition across 21 bone marrow samples analyzed by Binary-SPA, including controls (Ctrl-1 to Ctrl-4), monoclonal gammopathy of undetermined significance (MGUS-1 to MGUS-2), smoldering multiple myeloma (SM-1 to SM-5), and multiple myeloma (MM-1 to MM-10). Stacked bars represent the proportion of each cell type among all annotated cells. **b**, Plasma cell percentages detected by SingleR across disease stages. Each dot represents individual samples. Statistical significance compared to control was assessed by the Student’s t*-*test; ns = not significant. **c**, Plasma cell percentages detected by Binary-SPA across disease stages as in (b). ** p < 0.01, *** p < 0.001. **d-e**, Comparison of plasma cells annotated by Binary-SPA with CD138 immunohistochemistry staining^11^ in corresponding tissue sections. Plasma cells identified by Binary-SPA exhibit a spatial distribution consistent with that of CD138 staining. **f**, Correlation between computational annotations and clinical bone marrow plasma cell counts. Left: SingleR-derived plasma cell percentages versus clinical counts (Pearson r = 0.731, p = 0.000167). Right: Binary-SPA-derived plasma cell percentages versus clinical counts (Pearson r = 0.894, p < 1e-04). Each point represents one patient sample, colored by disease stage. **g**, Steiger’s test for comparing two dependent Pearson correlations sharing a common clinical reference variable. ** p < 0.01.

Pearson correlation analysis comparing annotated plasma cell percentages against clinical manual counts further demonstrated Binary-SPA’s superior accuracy: Binary-SPA achieved r = 0.894 (p < 1 × 10⁻⁴) compared with r = 0.731 (p = 1.67 × 10⁻⁴) for SingleR (**Fig. 5f**). A Steiger test confirmed that Binary-SPA’s correlation was significantly higher (**Fig. 5g**). These results demonstrate that Binary-SPA captures biologically and clinically meaningful variation in complex bone marrow samples where reference-dependent methods underperform.

### Binary-SPA enables accurate annotation of clinically archived bone marrow specimens

EDTA-based decalcification of the bone marrow core biopsy preserves RNA integrity but is not routinely performed in clinical practice. Most archival bone marrow specimens undergo harsh acid-based decalcification that causes substantial RNA degradation, making them unsuitable for spatial transcriptomics. Bone marrow clot biopsies, prepared from aspirate material that clots during collection, are standard components of clinical bone marrow evaluation and bypass decalcification entirely. Their routine availability in pathology archives makes them an attractive alternative for spatial transcriptomics analysis.

To evaluate this approach, we performed Xenium 5K analysis on a normal bone marrow clot biopsy sample and applied Binary-SPA for cell annotation (**Fig. 6a**). We subsequently performed cell type annotation using the same benchmarking methods evaluated above. For label-transfer–based methods, we used a published human bone marrow scRNA-seq reference dataset^25^. Binary-SPA achieved 100% annotation coverage and recovered all 19 predefined cell types. The label transfer method SELINA annotated fewer than 1% of cells and was excluded from further analysis. Marker-based methods TACIT and ScType achieved only 52.7% and 70% coverage, identifying 16 and 10 cell types, respectively (**Fig. 6b-d, Extended Data Table 6**).

**Fig. 6.**
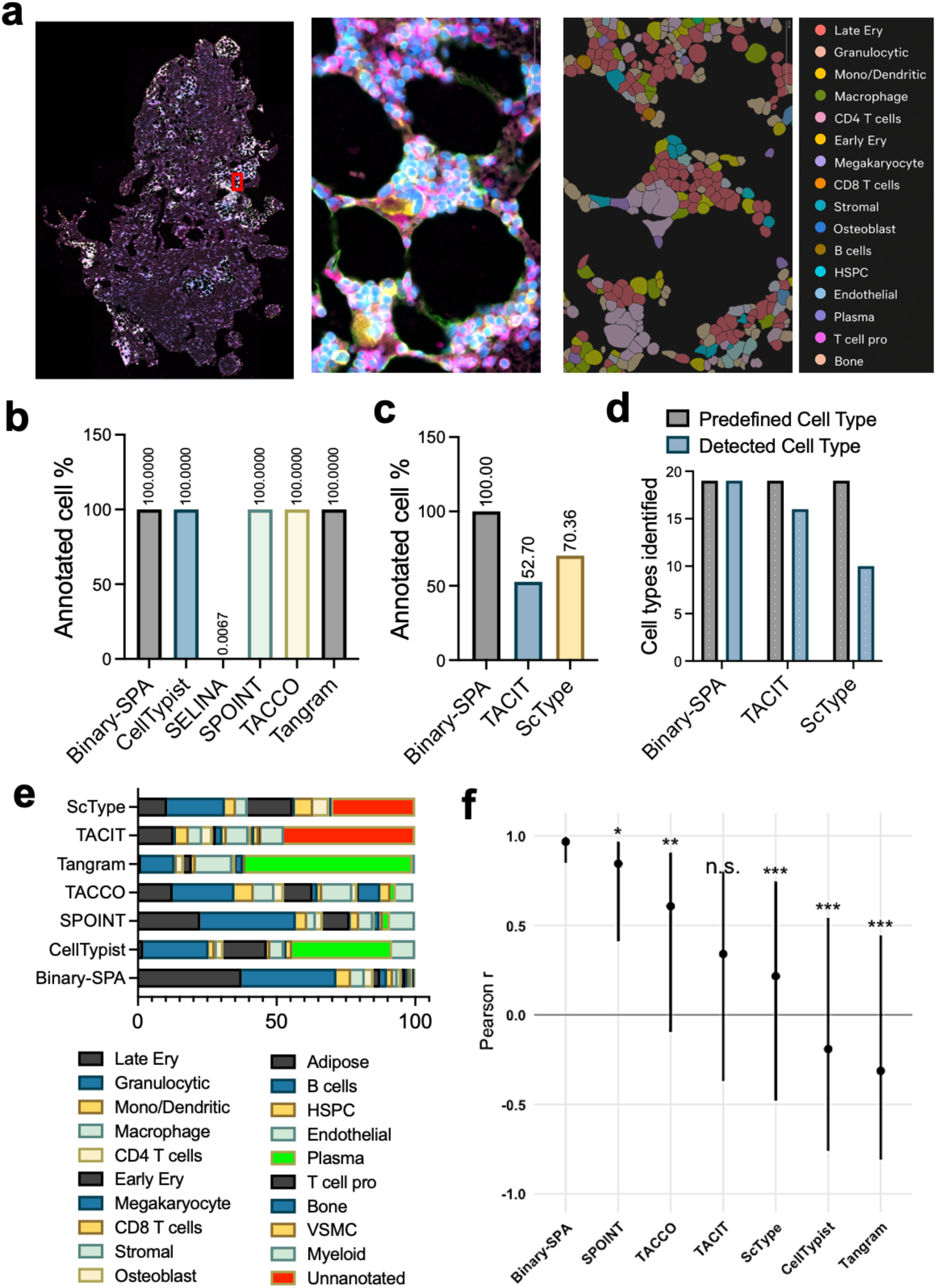
Binary-SPA enables accurate annotation of clinically archived bone marrow specimens. **a**, Representative Binary-SPA annotation of a normal bone marrow clot biopsy. Left: full-sample spatial data with a red box indicating the enlarged region. Middle: Xenium immunofluorescence signals showing DAPI-labeled nuclei, cell boundaries (red outlines), RNA transcripts (yellow), and protein signals (green) in the enlarged region. Right: Binary-SPA cell type annotations with corresponding color legend. **b**, Annotation coverage comparison between Binary-SPA and label transfer-based methods. SELINA showed an annotation rate of less than 1% and was excluded from downstream benchmarking analyses. **c**, Annotation coverage comparison between Binary-SPA and marker-based annotation methods. **d**, Cell type retrieval comparison across marker-based annotation approaches. **e**, Cell type composition across annotation methods. Each horizontal bar represents one method; colors indicate cell type proportions. **f**, Correlation of cell type proportions with Lunaphore COMET protein-based measurements. Points indicate the correlation coefficient (r), and vertical lines represent the 95% confidence intervals. Statistical significance was determined using Steiger’s test for overlapping correlations, comparing each method to Binary-SPA. Significance levels are denoted as ns, not significant; * p < 0.05; ** p < 0.01; *** p < 0.001.

Cell type proportion analysis revealed that Binary-SPA annotated the majority of cells as late erythroid cells and granulocytes, consistent with expected normal bone marrow composition. CellTypist and Tangram, despite strong performance with matched references in prior analyses, assigned disproportionately high fractions of cells as plasma cells at levels typically seen in plasma cell neoplasms rather than normal bone marrow (**Fig. 6e**).

To establish ground truth for quantitative benchmarking, we performed Lunaphore COMET multiplexed protein imaging on the same specimen and annotated cells using TACIT, which has demonstrated effectiveness for protein-based annotation^17^. Pearson correlation analysis comparing transcriptomics-derived annotations against COMET-derived cell type proportions revealed that Binary-SPA achieved r = 0.968, indicating near-perfect concordance with protein-based ground truth (**Fig. 6f**, **Extended Data Fig. 5, and Table 7**). All other methods exhibited substantially lower correlations with broader confidence intervals, and Steiger’s tests confirmed Binary-SPA’s significantly superior performance.

These findings demonstrate that Binary-SPA maintains high accuracy in archival specimens where reference-dependent methods fail due to a lack of appropriately matched reference data, a common limitation in clinical and translational research settings.

## Discussion

High-resolution spatial transcriptomics has emerged as a transformative technology with the potential to reshape both research and clinical practice. However, accurate cell type annotation remains a major bottleneck. Label transfer methods can achieve high accuracy when well-matched scRNA-seq references are available, but their performance degrades substantially when references are mismatched or derived from independent studies, a common scenario for archived clinical specimens. Marker-based methods eliminate reference dependency but often suffer from incomplete annotation coverage and failure to recover rare cell populations.

These limitations of marker-based approaches largely stem from reliance on expression levels to infer cell identity, which introduces two fundamental problems. First, it assumes RNA abundance directly reflects protein levels, yet bulk analyses indicate that mRNA explains only ∼40% of protein-level variation^26–28^. Moreover, transcription at the single-cell level is stochastic, occurring in burst-like ON/OFF states that generate substantial cell-to-cell variability in RNA abundance^28,29^. Second, highly expressed individual markers can disproportionately dominate cell-type scoring, creating AND/OR artifacts in which cells expressing a single abundant marker receive higher scores than those expressing multiple biologically relevant markers. In practice, the presence of multiple markers is often more informative for defining cell identity than the high expression of a single gene.

Binary-SPA directly addresses these issues through two key innovations. First, by converting marker gene expression into binary states (detected vs. non-detected), the resulting cell type score (CTS) depends on the number of positive markers rather than their absolute expression levels. This design prioritizes combinatorial marker expression and is particularly important for identifying cell types defined by multiple co-expressed markers. Second, the ΔCTS metric, the difference between the highest and second-highest normalized scores, enables robust discrimination even when marker detection is incomplete^28,30–32^, providing a self-adaptive mechanism that accommodates differences in platform sensitivity, sequencing depth, and sample quality.

A fundamental distinction of Binary-SPA is its cell-by-cell annotation strategy rather than cluster-based assignment. Conventional clustering approaches group cells based on global transcriptomic similarity using highly variable genes and dimensionality reduction^33^. While computationally efficient, this logic differs fundamentally from traditional cell typing, which defines identity based on a limited set of marker proteins. We observed that cells of the same biological type often distribute across multiple clusters, while individual clusters may contain multiple cell types. Binary-SPA bridges this gap by evaluating each cell individually against marker-defined criteria, producing annotations that align with classical cell classification frameworks and are more readily interpretable in practice.

The SPA component addresses a critical challenge: many cells lack detectable expression of sufficient markers due to technical limitations in sensitivity or biological variability. We reasoned that although these cells cannot be confidently annotated solely by markers, they retain global transcriptomic signatures characteristic of their true cell type. The self-referencing strategy exploits this principle by using high-confidence cells identified in the Binary step as internal anchors for label transfer. Because anchor cells share identical technical artifacts, batch effects, and biological conditions with the cells being annotated, label transfer operates within a homogeneous transcriptomic space, eliminating the domain shift that degrades performance when using external references. This design combines the interpretability of marker-based methods with the coverage of label transfer approaches.

We validated Binary-SPA across multiple platforms, preservation methods, and tissue types, including bone marrow, a particularly challenging tissue type. Bone marrow poses unique challenges for spatial transcriptomics due to RNA degradation caused by decalcification and the continuous developmental states of hematopoietic cells. Binary-SPA reliably identified cell populations in both EDTA-decalcified core biopsies and clot biopsies that bypassed decalcification entirely, capturing clinically meaningful differences across disease stages. Orthogonal validation using Lunaphore COMET protein imaging showed strong concordance between Binary-SPA annotations and protein-based cell identities (r = 0.968), demonstrating that the method achieves accuracy comparable to that of gold-standard proteomic approaches.

Because Binary-SPA relies on user-defined marker genes, optimal performance requires a comprehensive marker matrix that captures the expected cell populations in the tissue of interest. The quality control step at the beginning of the Binary-SPA workflow, in which unsupervised clustering is performed to identify unexpected populations, serves as a safeguard, ensuring that the marker matrix is updated before annotation proceeds.

Overall, Binary-SPA provides a robust framework for high-resolution spatial transcriptomics annotation that achieves 100% coverage while maintaining accuracy comparable to, or exceeding, that of label-transfer methods that depend on matched reference datasets. By eliminating the need for external references without sacrificing performance, Binary-SPA offers a scalable, practical solution for comprehensive spatial cell-type annotation across diverse research and clinical applications.

## Methods

### Datasets Used for Validation

The Xenium and Visium data for COAD, HCC, and OV were acquired from the SPATCH website (http://spatch.pku-genomics.org/) and the Genome Sequence Archive at the National Genomics Data Center under accession number HRA011129. The plasma cell neoplasia patient’s bone marrow Xenium data were acquired from the Gene Expression Omnibus database (accession number GSE299207).

### Binary-SPA pipeline structure

Prior to annotation, quality control is performed using unsupervised clustering to verify that no unexpected cell populations are present in the sample. Differentially expressed genes are examined across clusters to identify previously unannotated cell types or cells originating from other tissues (e.g., metastatic populations). If unexpected populations are detected, corresponding markers are added to the marker matrix before proceeding.

A user-defined marker matrix is then constructed with cell types as rows and marker genes as columns. For each cell type, marker genes expected to be positively expressed based on prior knowledge are assigned a value of 1, while all other entries are set to 0. These markers can correspond to classical cell-typing markers used in flow cytometry or immunofluorescence staining (exemplified in Extended Data Tables 1–5).

The pipeline begins by generating a cell-by-gene expression matrix. This can be achieved in two ways: (1) by directly reading spatial transcriptomics data using Seurat::Read10X or Seurat::Read10X_h5 to load the cell feature matrix from Xenium output, followed by extraction of the expression matrix using Seurat::GetAssayData; or (2) by importing a precomputed AnnData object (.h5ad) using anndataR::read_h5ad, from which the expression matrix is extracted for downstream analysis.In the resulting matrix, each row corresponds to a cell, each column corresponds to a gene, and each entry represents the expression level of a given gene in a given cell.

Because spatial transcriptomics platforms vary in gene coverage, some marker genes in the user-defined matrix may not be present in the dataset. To ensure compatibility, a platform-specific marker matrix (“marker matrix of use”) is generated by intersecting the user-defined marker genes with genes detected in the spatial dataset. Marker genes absent from the spatial data are removed, retaining only those that can be evaluated. This step enables automatic adaptation across different platforms without manual curation.

The cell-by-gene expression matrix is subset to include only marker genes present in the marker matrix of use, then binarized: detectable expression (value > 0) is assigned 1, and non-detectable expression is assigned 0. This converts marker gene expression into a simple on/off signal, prioritizing the presence of multiple markers over expression magnitude for any single marker.

Subsequently, a cell type score (CTS) is calculated by multiplying the marker matrix of use (cell types × marker genes) by the transposed binarized expression matrix (marker genes × cells), yielding a CTS matrix in which each cell receives a score for each predefined cell type. Then, the CTS matrix is normalized within each cell type, yielding a CTS value for each cell. For each cell, the difference between the highest and second-highest normalized CTS values is calculated to obtain the ΔCTS. A threshold is then defined (see below for details), and cells with ΔCTS values exceeding this threshold are classified as clear cells. These clear cells are subsequently used as reference cells for the SPA process.

In the SPA step, clear cells serve as the reference dataset for anchor-based label transfer onto unclear cells. Anchor-based label transfer is performed using Seurat::FindTransferAnchors() to identify anchor cell pairs, followed by Seurat::MapQuery() to project annotations from clear cells onto unclear cells. Because all cells originate from the same sample and share identical experimental conditions, batch effects are minimized, enabling accurate label transfer without external reference data. The predicted annotations are merged back into the original dataset, yielding a complete annotation of all cells.

### Cell type score (CTS) calculation

We computed a cell type score (CTS) based on binarized marker gene expression. Let

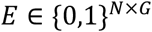

denote the binarized marker expression matrix, where *N* represents cells and *G* represents marker genes. An entry *E_ij_* = 1 indicates that marker gene *j*is detectably expressed in cell *i*, whereas *E_ij_* = 0 indicates non-detectable expression.

Let

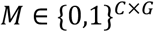

denote the marker matrix, where *C* represents cell types, and an entry *M_kj_* = 1 indicates that gene *j* is a marker for cell type *k*. To align matrix dimensions for multiplication, the binarized marker expression matrix *E* is transposed. The CTS matrix is then computed as:

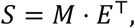

yielding

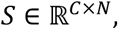

where each element

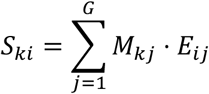

represents the CTS of cell *i* for cell type *k*.

### Calculation of ΔCTS

CTS were calculated for each cell using predefined marker gene sets for candidate cell types, as described above. Thus, each cell received a CTS for every candidate cell type.

For each cell, CTS values were normalized using min–max scaling so that all scores ranged from 0 to 1. The Delta Cell Type Score (ΔCTS) for each cell *i*was defined as:

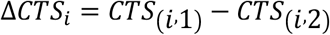

where *CTS*_(*i*,1)_ represents the highest normalized cell type score for cell *i*, and *CTS*_(*i*,2)_ represents the second-highest normalized cell type score for the same cell.

The ΔCTS threshold was determined based on tissue context and data quality. For consistency and comparison across samples in this study, a uniform ΔCTS cutoff of 0.15 was applied to define confident cell type assignments. Cells with ΔCTS ≥ 0.15 were classified as clear cells, whereas cells with ΔCTS < 0.15 were classified as unclear cells.

### Self-referenced projection annotation (SPA) using clear cells as reference

For self-referenced projection annotation (SPA), cells were split into clear and unclear groups based on the ΔCTS threshold as described above. Clear cells were used as the reference object, and unclear cells were used as the query object. Both datasets were processed in parallel using Seurat’s standard workflow: NormalizeData, FindVariableFeatures, ScaleData, and RunPCA (30 PCs).

Transfer anchors between the reference and query were identified using FindTransferAnchors with normalization.method = “LogNormalize”, reference.reduction = “pca”, and dims = 1:30. The query cells were then projected onto the reference using MapQuery, transferring cell type labels (refdata = label_col) and mapping the query into the reference PCA space and UMAP embedding (reference.reduction = “pca”, reduction.model = “umap”).

After projection and label transfer, the reference and query datasets were merged using the Seurat “merge” function to generate a unified object for downstream analysis.

### Marker sets for Binary-SPA annotation

For FFPE Xenium data from COAD and OV, the marker sets used for Binary-SPA annotation are provided in **Extended Data Table 1**. Marker sets for the HCC sample are listed in **Extended Data Table 2**. Marker sets for Visium fresh-frozen (FF) and FFPE COAD datasets are listed in **Extended Data Table 3**. Marker sets used for bone marrow datasets are summarized in **Extended Data Table 4**.

### Benchmarking against label transfer methods

All label transfer methods used matched scRNA-seq reference datasets provided in the original publications. These datasets were obtained from the SPATCH website (http://spatch.pku-genomics.org/)^11^.

#### Data loading and preprocessing

Spatial transcriptomics data were loaded from AnnData (adata.h5ad) objects. A matched scRNA-seq reference AnnData object was loaded for each sample and used as the annotation reference. The scRNA-seq reference cell type label was obtained from ref.obs[“major_annotation”].

#### CellTypist

CellTypist was applied using the scRNA-seq reference. A CellTypist model was trained on the reference AnnData with feature_selection=True, using the reference major cell type labels (major_annotation) as training labels. The trained model was then used to annotate Xenium cells.

#### Tangram

Tangram uses the reference dataset to project reference labels onto query cells. We first filter for overlapping genes using the intersection of the reference and Xenium gene sets. Next, both objects were subset to the shared gene space prior to mapping. Reference annotations were then projected onto Xenium cells using Tangram’s label projection function.

##### TACCO

TACCO was applied using the scRNA-seq dataset as the reference. The reference and Xenium query datasets were harmonized by intersecting on shared genes using numpy.intersect1d. Cell type annotation was then performed using the tacco.tl.annotate function.

##### SPOINT

SPOINT integrates transcriptomic similarity with spatial context to infer cell type identities. A scRNA-seq dataset served as the reference, and the model was trained using the reference-derived major cell type labels. Next, spatial deconvolution was performed using the deconv_spatial function to compute per-cell cell type prediction scores for the Xenium dataset. The annotated spatial AnnData was then retrieved using spoint_model.st_ad.

##### SELINA

SELINA was run in a virtual environment following the authors’ recommended setup^12^. Query Xenium data were preprocessed using *selina preprocess*. A scRNA-seq reference was used to train the model with *selina train*, and the trained model was applied to annotate the Xenium dataset using *selina predict*.

### Benchmarking against marker-based annotation methods

#### TACIT

Xenium data were preprocessed to generate a cell-by-gene matrix for TACIT annotation. The same marker gene set used in Binary-SPA was applied, and TACIT was executed with parameters r = 10 and p = 20.

#### ScType

Xenium spatial transcriptomics data were imported using the *readH5AD* function and converted into a Seurat object using *as.Seurat.* Cell type annotation was then performed using ScType with marker genes matched to those used in Binary-SPA to ensure consistency across methods. For COAD and OV datasets, marker genes listed in **Extended Data Table 4** were used with general labels. For HCC datasets, marker genes included both general markers and liver markers. For bone marrow datasets, marker genes listed in **Extended Data Table 6** were used.

### Binned Pearson Correlation Analysis

To quantify spatial concordance between transcriptomics-derived annotations and CODEX protein-based ground truth, binned Pearson correlation analysis was performed based on cell-type composition within spatial tiles.

Xenium cell centroid coordinates were partitioned into approximately 1,000 spatial tiles by defining bin boundaries along the x- and y-axes using coordinate quantiles. These boundaries were then applied to both Xenium and CODEX datasets to assign each cell a shared tile identifier based on spatial location, enabling direct comparison of cell-type composition at matched spatial positions. Cells falling outside defined tile boundaries were excluded from analysis.

Within each tile, cell-type proportions were calculated independently for each annotation method and for CODEX. Pearson correlation coefficients were then computed between the cell-type proportion vector from each annotation method and the corresponding CODEX-derived proportion vector for that tile. Per-tile correlation values were aggregated across all tiles to generate distributions for statistical comparison between methods. Wilcoxon signed-rank tests were used to assess whether correlation distributions differed significantly between Binary-SPA and alternative annotation approaches.

### Human samples collection

Human bone marrow clot section samples were obtained from leftover diagnostic specimens at Northwestern Memorial Hospital. The study protocol was approved by Northwestern University’s institutional review board (IRB ID: STU00217116). Human sample embedding and sectioning were performed at Northwestern University’s Pathology Core Facility.

### Xenium spatial transcriptomic analysis of bone marrow clot biopsies

Xenium transcriptome spatial RNA sequencing platforms for humans were purchased from 10X Genomics (Pleasanton, USA). A 5-µm bone marrow clot section tissue curl was used to assess RNA quality. Samples that passed quality control were subsequently selected for Xenium analysis. All analyses were performed at the Northwestern University NUSeq Core Facility. Briefly, spatial transcriptomic analysis was performed using the Xenium Prime 5K gene expression assay. FFPE sections were mounted on Xenium slides and prepared according to the manufacturer’s tissue preparation workflow for deparaffinization, decrosslinking, probe hybridization, and amplification. Prepared slides were run on the Xenium Analyzer, and the resulting data were processed using the Xenium Onboard Analysis.

### Spatial omics analysis using Lunaphore COMET

A matching bone marrow clot section tissue block from the sample analyzed by Xenium was sectioned for multiplexed protein analysis. Briefly, 5-µm sections were cut and mounted onto high-adhesive glass slides (TOMO). Multiplexed immunofluorescence staining was performed using a panel of 14 antibodies. The antibody panel included c-Kit (Abcam, ab317843), CD34 (Abcam, ab81289), CD61 (Cell Marque, 161M-14), E-cadherin (Cell Signaling Technology, 14472), CD71 (Cell Marque, 171M-94), hemoglobin (Invitrogen, MA1-10806), myeloperoxidase (MPO; Abcam, ab9535), CD163 (Cell Marque, 163M-14), CD68 (Invitrogen, MA5-13324), CD3 (Cell Marque, 103R-95), CD4 (Abcam, ab133616), CD8 (Bio-Rad, MCA1817), CD20 (Cell Marque, 120M-85), and CD45 (Cell Marque, 145M-9).

Prepared slides were processed by the University of Alabama at Birmingham (UAB) Flow Cytometry and Single Cell Core Facility for downstream analysis using the Lunaphore COMET platform (Tolochenaz, Switzerland). COMET staining, imaging, and signal extraction were performed according to the manufacturer’s recommended protocols at the UAB core facility.

### COMET data processing and cell annotation

Multiplexed imaging data generated by the Lunaphore COMET platform were processed using QuPath for visualization and cell segmentation. Nuclear segmentation was performed based on DAPI staining, with segmentation masks expanded by 5 µm to approximate whole-cell boundaries. Fluorescence intensity measurements for each marker were extracted at the single-cell level and exported for downstream analysis.

To reduce background noise, fluorescence signals were corrected by subtracting the nonspecific signal derived from the corresponding secondary antibody control. The resulting cell/feature matrix was subsequently imported into TACIT for cell type annotation and downstream analysis using the marker set listed in **Extended Data Table 6**.

## Data availability

The COAD, OV, and HCC Xenium, scRNA-seq, and CODEX datasets were downloaded from the SPATCH database (https://spatch.pku-genomics.org/#/homepage). The multiple myeloma (MM) dataset was obtained from the Gene Expression Omnibus (GEO) under accession number GSE299207. The bone marrow clot biopsy Xenium dataset generated in this study has been deposited in GEO under accession number GSE322974. The Lunaphore COMET dataset has been deposited in Figshare under DOI: 10.6084/m9.figshare.31502971.

## Code availability

The code for Binary-SPA is available on GitHub at https://github.com/HonghaoNU/BinarySPA. For information regarding commercial use or licensing inquiries, please refer to the LICENSE file in the repository or contact the corresponding author.

## Acknowledgments

We thank the Mouse Histology and Phenotyping Laboratory, the Pathology Core Facility, and the NUSeq Core Facility at Northwestern Feinberg School of Medicine for their assistance with tissue preparations and assays. This work was supported by the National Institute of Diabetes and Digestive and Kidney Disease (NIDDK) grant R01-DK124220 (P.J.) and R01-DK138205 (P.J.). National Heart, Lung, and Blood Institute (NHLBI) grant R01-HL148012 (P.J.), R01-HL169507 (P.J.), and R01-HL150729 (P.J.). H.B. is a recipient of the F32 Ruth L. Kirschstein Postdoctoral Individual National Research Service Award (F32-HL170648).

## Author contributions

H.B., W.C., P.W., K.R., I.A., and E.L. performed the experiments and interpreted data. C.W. and M.J.S. performed the sequencing experiments. J.M-C. interpreted data. H.B. and P.J. designed the experiments, interpreted data, and wrote the manuscript.

## Ethics declarations

The authors declare no competing interests related to this work.

