## Extended Figures for "Binary-SPA: A Reference-Free Method for Cell Annotation in High-Resolution Spatial Transcriptomics"

### Extended Data
### Figures and Figure Legends

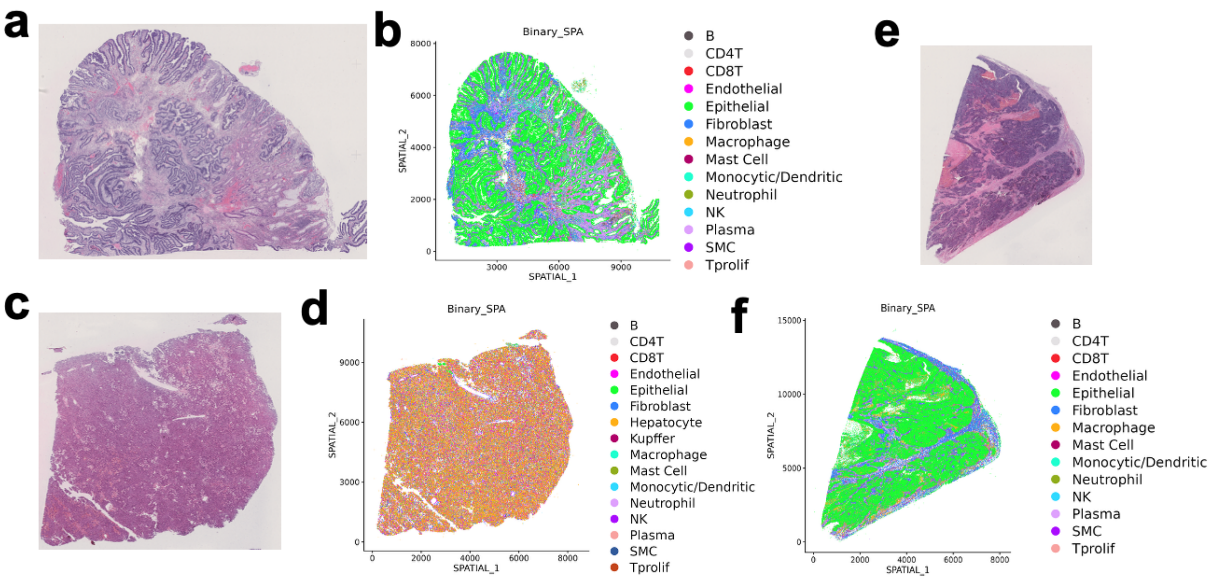


**Extended Data Fig. 1. Binary-SPA annotation of Xenium 5k data across three tumor types with corresponding histology. a-b,** H&E-stained colon adenocarcinoma (COAD) tissue section and corresponding Binary-SPA annotation map showing spatial distribution of cell types. **c-d**, H&E-stained hepatocellular carcinoma (HCC) tissue section and corresponding Binary-SPA annotation map. Note the inclusion of hepatocytes (orange) specific to liver tissue. **e-f**, H&E-stained ovarian cancer (OV) tissue section and corresponding Binary-SPA annotation map.

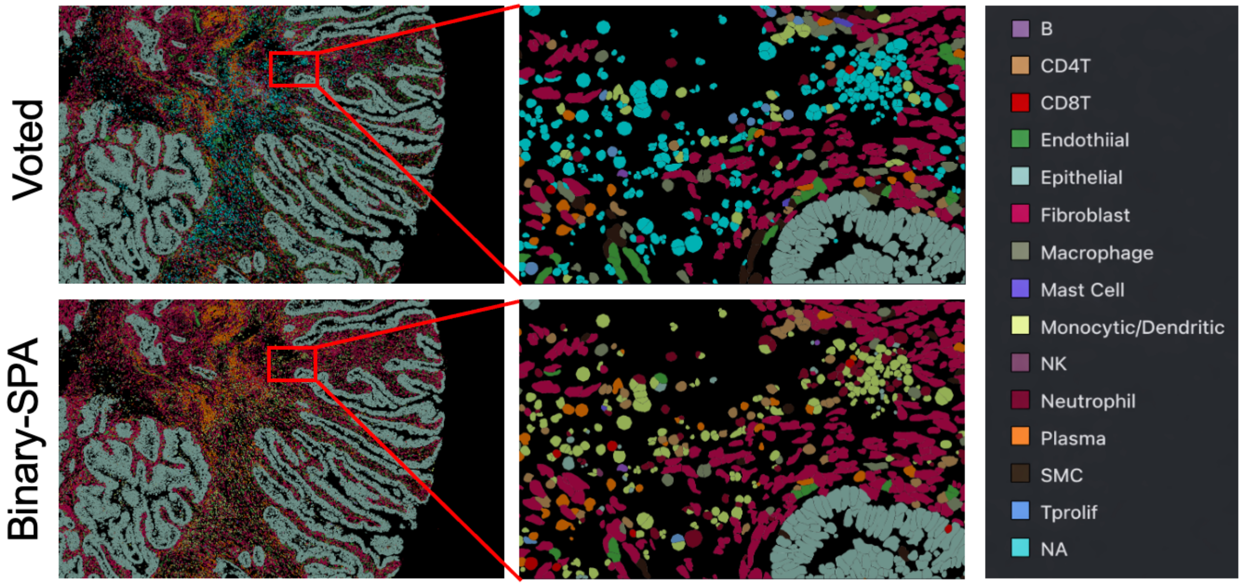


**Extended Data Fig. 2. Binary-SPA successfully annotates cells that remained unclassified by the original voting-based method.** Representative spatial plot from COAD Xenium data showing cells that were labeled as "NA" (unannotated) by the original voting-based consensus method. These cells, which failed to reach annotation consensus across five reference-dependent methods, were successfully assigned cell type identities by Binary-SPA. Cell types are color-coded as indicated in the legend.

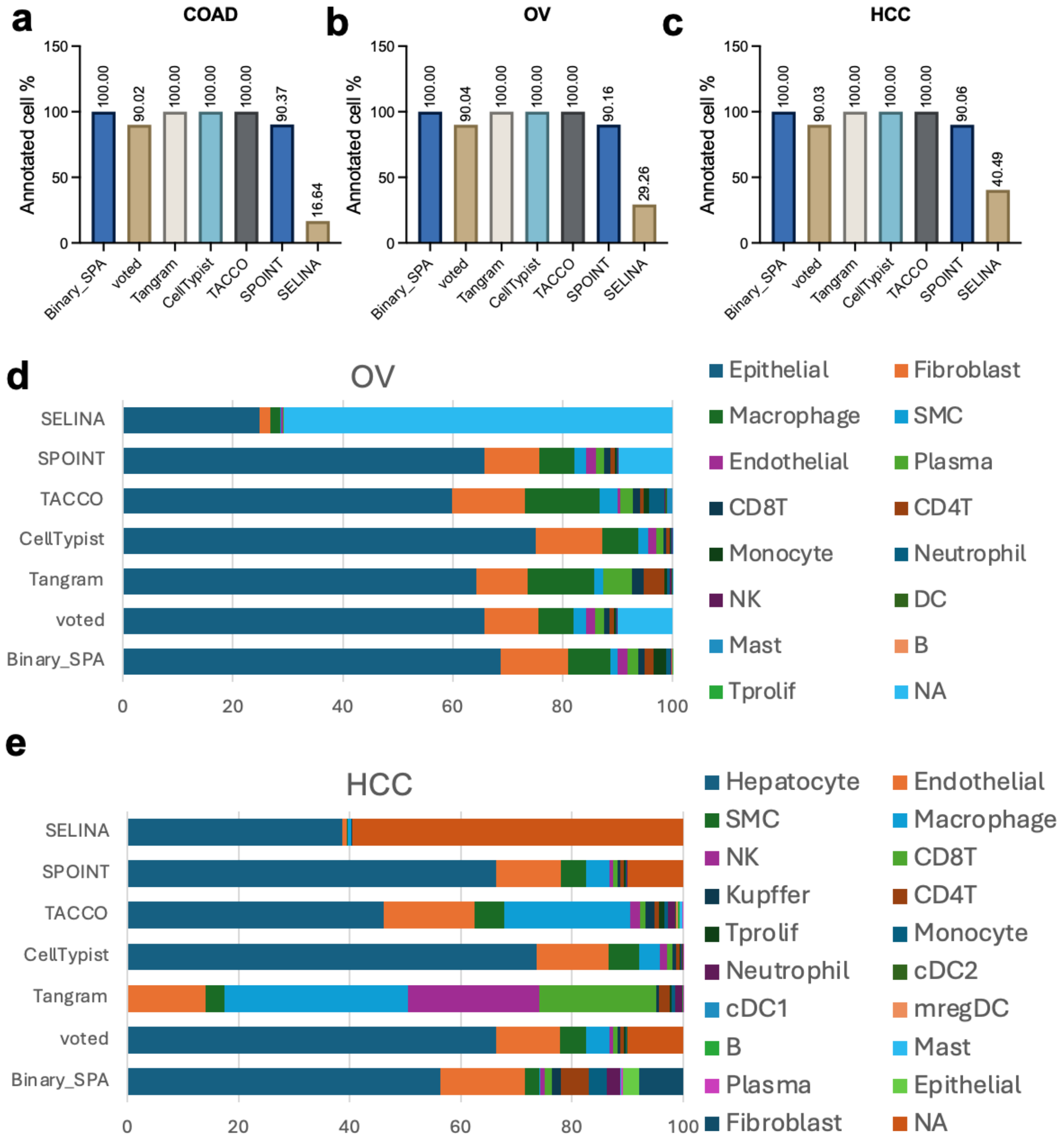


**Extended Data Fig. 3. Annotation coverage and cell type proportion comparison between Binary-SPA and label transfer–based methods. a-c**, Percentage of successfully annotated cells by each method in COAD (a), OV (b), and HCC (c) Xenium datasets. **d**, Cell type proportion comparison in OV Xenium samples between Binary-SPA and five label transfer-based methods. **e**, Cell type proportion comparison in HCC Xenium samples between Binary-SPA and five label transfer-based methods.

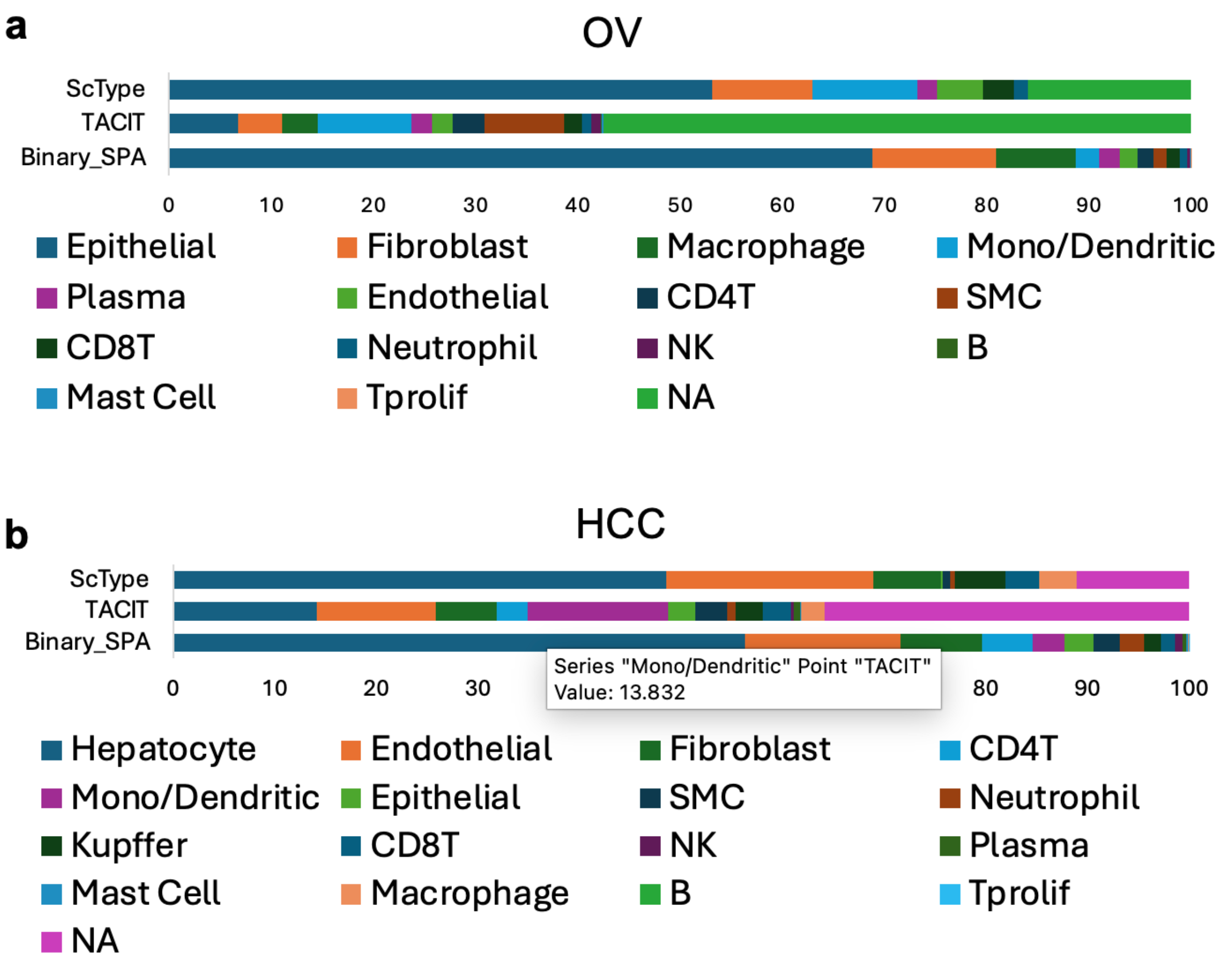


**Extended Data Fig. 4. Cell type proportion comparison between Binary-SPA and marker-based annotation methods. a**, Cell type proportions in OV Xenium samples annotated by Binary-SPA, TACIT, and ScType using identical marker gene sets. **b**, Cell type proportions in HCC Xenium samples annotated by the same three methods.

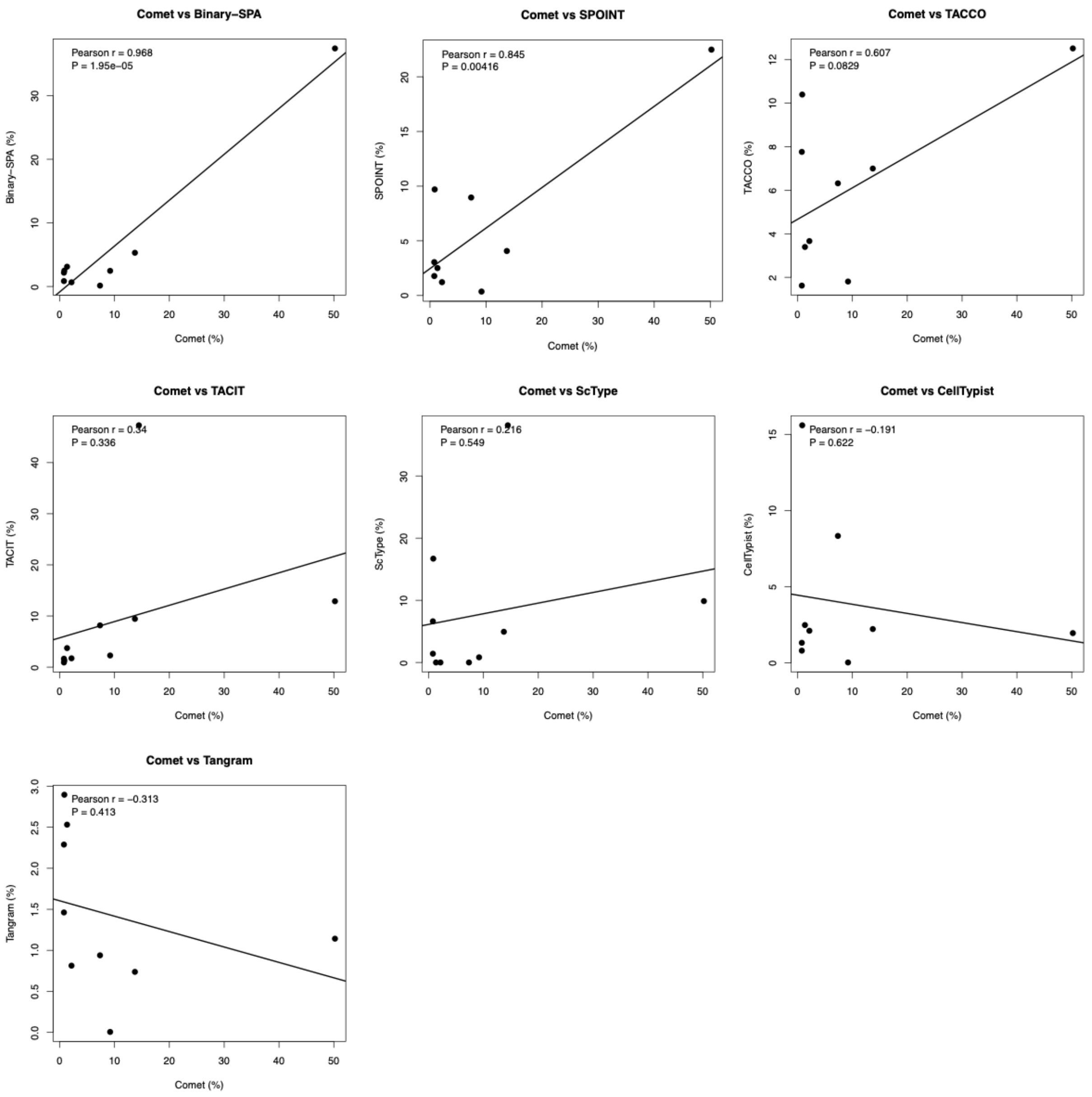


**Extended Data Fig. 5. Concordance between annotation methods and Lunaphore COMET protein-based ground truth in bone marrow clot biopsy.** Pearson correlation analysis comparing cell type proportions derived from each transcriptomics-based annotation method against corresponding proportions from Lunaphore COMET multiplexed protein imaging. Error bars represent 95% confidence intervals.

**Extended Data Table Legends**

**Extended Data Table 1.** Marker gene matrix used for Binary-SPA analysis of COAD and OV Xenium datasets.

**Extended Data Table 2**. Marker gene matrix used for Binary-SPA analysis of HCC Xenium dataset.

**Extended Data Table 3**. Marker gene matrix used for Binary-SPA analysis of COAD Visium HD dataset.

**Extended Data Table 4**. Marker gene matrix used for Binary-SPA analysis of all Bone marrow Xenium datasets.

**Extended Data Table 5**. Marker gene matrix used for ScType analysis of COAD, OV and HCC Xenium datasets. General marker sets were used for COAD and OV, and the liver-specific marker set was used for HCC.

**Extended Data Table 6**. Marker gene matrix used for ScType analysis of bone marrow Xenium datasets.

**Extended Data Table 7**. Marker gene matrix used for TACIT analysis of the bone marrow clot biopsy Lunaphore COMET dataset.
